# A Cdc42-Borg4-Septin 7 axis regulates HSCs polarity and function

**DOI:** 10.1101/2021.03.17.435817

**Authors:** Ravinder Kandi, Katharina Senger, Ani Grigoryan, Karin Soller, Vadim Sakk, Tanja Schuster, Karina Eiwen, Manoj B. Menon, Matthias Gaestel, Yi Zheng, Maria Carolina Florian, Hartmut Geiger

## Abstract

Aging of hematopoietic stem cells (HSCs) is caused by an elevated activity of the small RhoGTPase Cdc42 and an apolar distribution of proteins. Mechanisms by which Cdc42 activity controls polarity of HSCs are not known. Binder of RhoGTPases proteins (borgs) are known effector proteins of Cdc42 that are able to regulate the cytoskeletal septin network. Here we show that Cdc42 interacts with borg4, which in turn interacts with septin 7 to regulate the polar distribution of Cdc42, borg4 and septin 7 within HSCs. Genetic deletion of either borg4 or septin 7 in HSCs resulted in a reduced frequency of HSCs polar for Cdc42 or borg4 or septin 7 and a reduced engraftment potential and decreased lymphoid-primed multipotent progenitors (LMPPs) frequency in the bone marrow. In aggregation our data identify a Cdc42-borg4-septin 7 axis to be essential for maintenance of polarity within HSCs and for HSC function and provide rationale for further investigating the role of borgs and septins for the regulation of compartmentalization within stem cells.

**Graphical Abstract:** 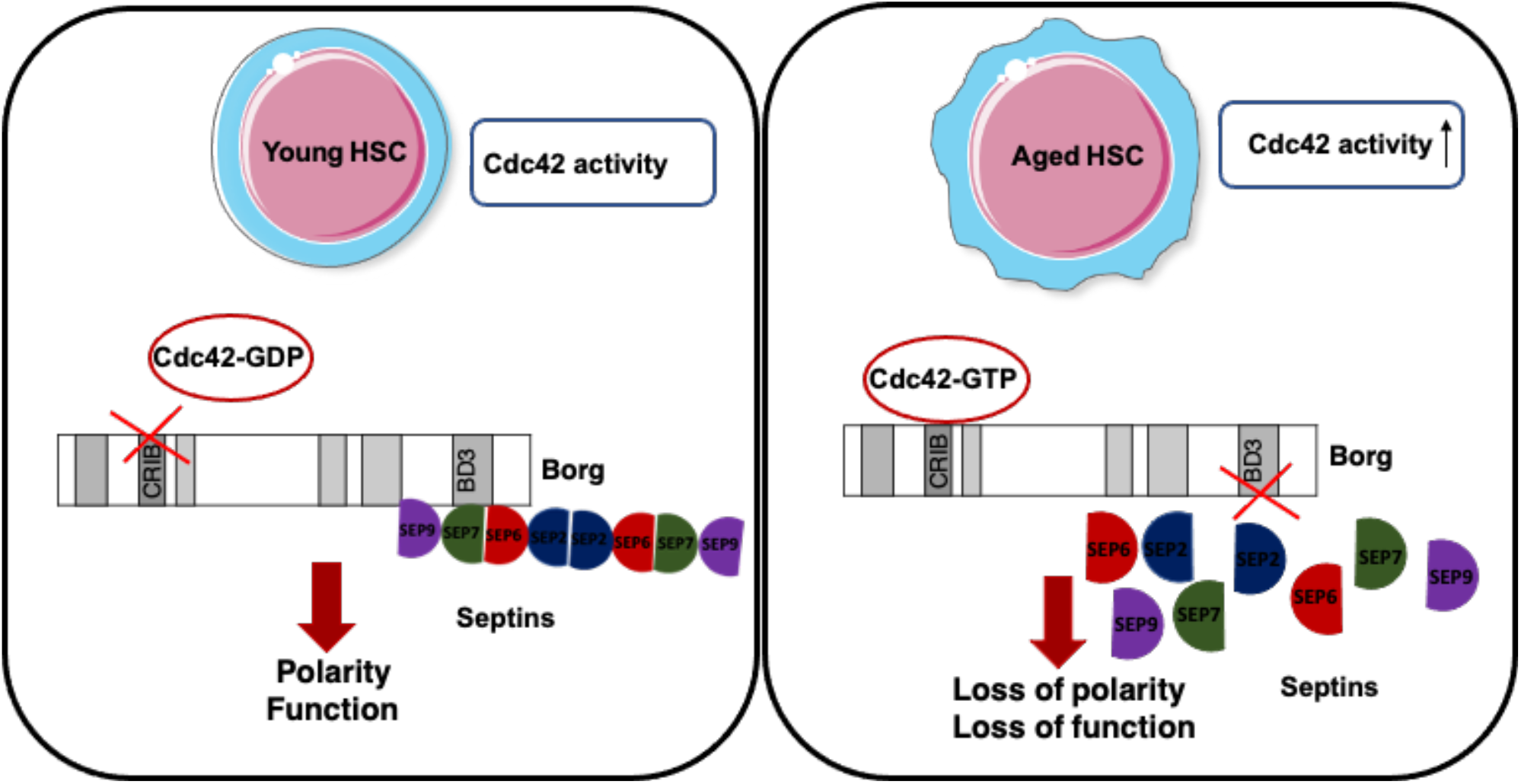

## Introduction

The function of hematopoietic stem cells (HSCs) decreases upon aging. This contributes, among others, to aging associated immune remodelling (AAIR) with a reduced production of lymphoid cells (B-cells and T-cells) and erythroid cells in bone marrow (BM) as well as an increased myeloid output linked to aging-related myeloid malignancies (Akunuru & Geiger, 2016; Geiger *et al*, 2013; Snoeck, 2013; Beerman *et al*, 2010; Rossi *et al*, 2008). A prominent aging-related phenotype of aged HSCs is a reduced frequency of HSCs with a polar distribution of polarity proteins like Cdc42 or the cytoskeletal protein tubulin or the epigenetic marker histone 4 acetylated on lysine 16 (H4K16ac) (Florian *et al*, 2012). Polarity within HSCs is tightly linked to the mode of symmetrical or asymmetrical division. Young (polar) HSCs divide more asymmetrically, while aged (apolar) HSCs divide more symmetrically (Florian *et al*, 2018).The aging-related changes in HSCs and haematopoiesis are caused by both cell intrinsic alterations in HSCs and extrinsic BM niche factors (Geiger *et al*, 2007; Kamminga & de Haan, 2006; Wang *et al*, 2011; Guidi *et al*, 2017; Saçma *et al*, 2019; Mejia-Ramirez & Florian, 2020). An increase in the activity of the small RhoGTPase Cdc42 in aged HSCs causes the reduced frequency of polar HSCs upon aging and is also causative for HSC aging (Florian *et al*, 2012; Leins *et al*, 2018; Grigoryan *et al*, 2018; Liu *et al*, 2019; Amoah *et al*, 2021). Pharmacological attenuation of the aging-related increase in the activity of Cdc42 by a specific small molecule inhibitor of Cdc42 activity termed CASIN resets the frequency of polar HSCs back to level reported for young HSCs and rejuvenates the function of chronologically aged HSC. Cdc42 activity is thus a critical regulator of HSC polarity and aging (Florian *et al*, 2020).While there is information on polarity regulation pathways orchestrated by Cdc42 activity in yeast (Okada *et al*, 2013; Chollet *et al*, 2020), the mechanisms by which Cdc42 controls polarity and especially how elevated activity of Cdc42 result in loss of polarity in HSCs are not understood.

Ectopic expression of either a constitutively active form of Cdc42 or a dominant negative form of Cdc42 leads to redistribution of the borg family of Cdc42 effector proteins (also called Cdc42ep1-5) and subsequent loss of septin filaments in cell lines (Farrugia & Calvo, 2017; Joberty *et al*, 2001). Septins are GTP-binding proteins that interact in stable stoichiometry within each other to form filaments that bind to the cell membrane, actin filaments and microtubules. Septins are regarded as the fourth component of the cytoskeleton (Mostowy & Cossart, 2012). In yeast, septins are recruited to the site of polarization by active Cdc42 while at the same time inhibiting Cdc42 activity in a negative feedback loop (Okada *et al*, 2013) In budding yeast gic-1, a functional homologue of mammalian Borg proteins, binds to cdc42-GTP which in turn leads to dissociation of gic1 from septin filaments (Sadian *et al*, 2013, 1; Brown *et al*, 1997). Septins have gained recently more attention, also with respect to the hematopoietic system, like we and others have reported distinct roles for septin 6 (Senger *et al*, 2017) and septin 1(Ni *et al*, 2019) in hematopoiesis, while the borg family of Cdc42 effector proteins still remains largely uncharacterized, particularly within the hematopoietic system. Here, we demonstrate that a Cdc42-borg4-septin 7 axis regulates the intracellular distribution of polarity proteins in HSCs in response to changes in Cdc42 activity. Consequently, either borg4 or septin 7 are critical for HSCs function upon stress. Our results provide a novel scientific rationale for a role of borgs and septins in the compartmentalization of components within stem cells likely to be essential for stem cell function.

## Results

### Borg4 and Septin 7 show a polar distribution in HSCs which is regulated by Cdc42 activity

There are 5 known distinct binders of Rho GTPases proteins (Borgs), also known as Cdc42ep4 1-5, which can serve as Cdc42 effector proteins. Borgs are able to bind septins via a conserved BD3 domain and interact with Cdc42 through a Cdc42/Rac interactive binding (CRIB) motif. Borgs plays an important role in cytoskeletal rearrangement. They also regulate cell shape, filopodia formation and cell migration and which are controlled by specific interaction with active Cdc42-GTP(Farrugia & Calvo, 2016). We first determined the level of expression of borgs in HSCs. Borg 2-4 were expressed in LT-HSCs (Lin,c-Kit^+^,Sca-1^+^,flk2^-^,CD34^-^ cells) (Fig 1A), whereas borg 1 and 5 were absent or below the level of detection (data not shown). Expression of borg 2 and 3 was increased and borg4 was decreased in aged (elevated level of Cdc42 activity) compared to young LT-HSCs. There are 13 distinct mammalian septins. Septin 2,6,7,8,9,10 and 11 are thought to be ubiquitously expressed, while expression of septin 1,3,4,5,12 or 14 is usually restricted to distinct tissues or cell types. Septin 1,2,6,7,8,9 and 11 were indeed expressed in LT-HSC, while septin 3,4,5,10,12 and 14 were absent or below the level of detection (data not shown). The expression of septin 1,2,6 and 7 was decreased in aged LT-HSCs, while expression of septin 8 was increased upon aging (Fig 1B). The level of expression of septin 9 and 11 was similar in young and aged LT-HSCs. In summary a distinct set of borgs and septins are expressed in LT-HSCs, and most borgs and septins that are expressed in HSCs show changes in expression in aged compared to young HSCs.

**Figure 1.**
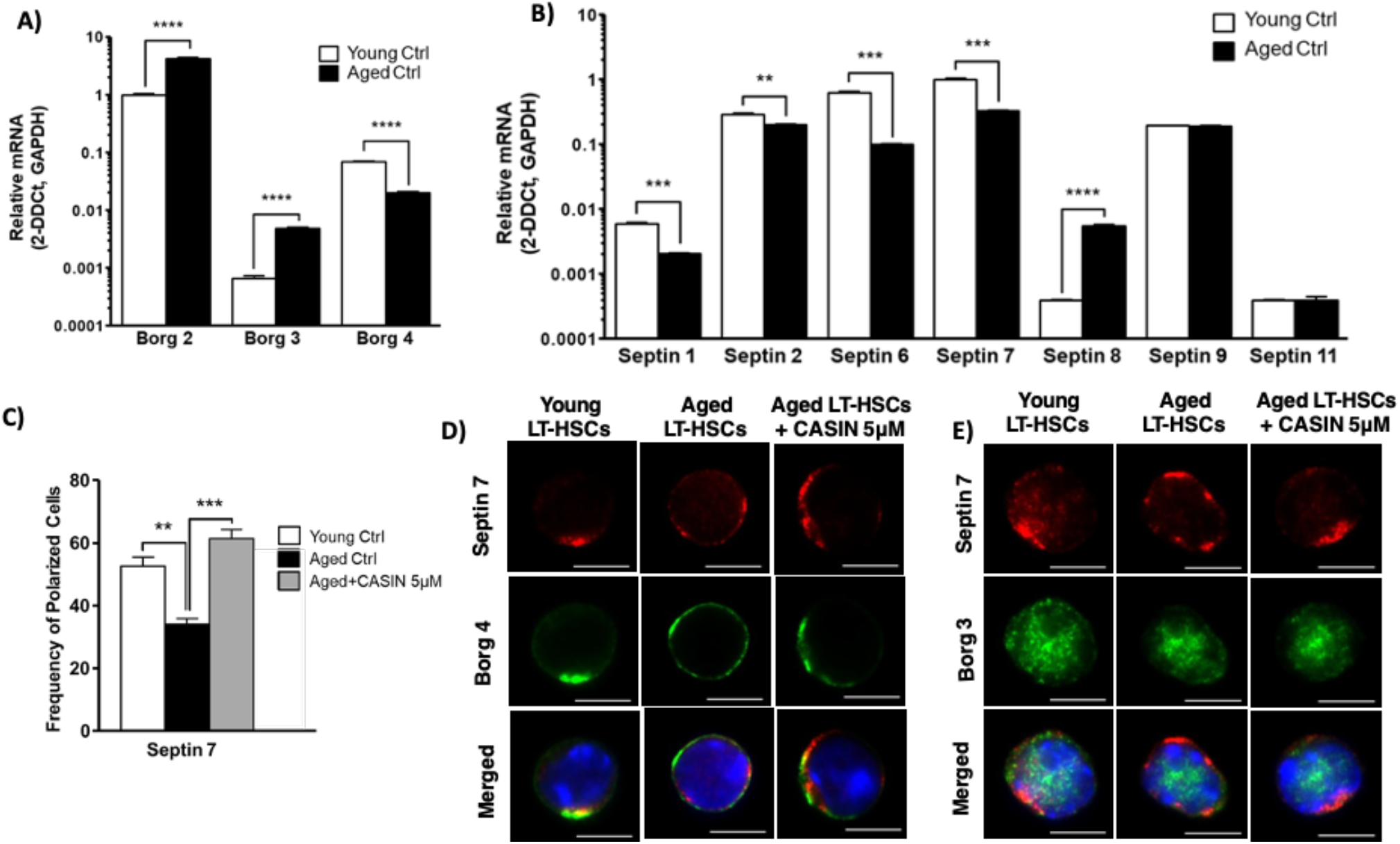
Borg4 and septin 7 show a polar distribution in HSCs which is regulated by Cdc42 activity. (A&B) Level of expression of borgs and septins in young and aged LT-HSCs. mRNA levels of borgs and septins were normalized to the level of expression of GAPDH and borgs are shown relative to the level of Borg 2 expression in young LT-HSCs. Septins are shown relative to the level of septin 7 expression in young LT-HSCs (n=3). Shown are mean±SEM, unpaired t-test, **P<0.01, ***P<0.001, ****P<0.0001. (C) Percentage of cells polarized for septin 7 among young, aged and aged LT-HSCs treated with CASIN (5µM). At least 3 biological repeats, at least 50 cells were scored per sample. Shown are mean±SEM, unpaired t-test, **P<0.01, ***P<0.001. (D&E) Representative immunofluorescence microscopy images of the distribution of septin 7 (red) or borg3 or borg4 (green) in young, aged and aged LT-HSCs treated with CASIN (5µM). Nuclei are stained with DAPI (blue), scale bar= 5µm.

Septins function in a physiological manner to align with scaffolds (tubulin, actin and vimentin) to allow for compartmentalization. In yeast, septins control the diffusion of specific proteins between mother and bud cells, regulating cellular aging (Takizawa *et al*, 2000; Shcheprova *et al*, 2008). We therefore investigated whether the spatial distribution of septin proteins is distinct in young and aged LT-HSCs using immunofluorescence analyses, and if so, whether changes in the distribution are a consequence of the elevated level of Cdc42 activity in aged LT-HSCs. The core septin filament structure consists of septins 2, 6 and 7 (Low & Macara, 2006; Sirajuddin *et al*, 2007). We therefore focused first on the localization of septin 2, 6 and 7 in HSCs. The distribution of septin 7 was polarized in about 50% of young but only in about 30% of aged LT-HSCs (Fig 1C and D), while only a very low frequency of LT-HSC (young or aged) were polar for either septin 2 or septin 6 (Fig S1K-M).

Aged HSCs possess an elevated activity of Cdc42 (more Cdc42-GTP) in comparison to young HSCs. The level of Cdc42 activity has been shown to control the spatial distribution of septins via borg effector proteins in both yeast and mammalian cell lines (Joberty *et al*, 2001). The distribution of septins in HSCs might therefore be regulated by the interaction of septins with the borg family of Cdc42 effector proteins (Sheffield *et al*, 2003; Farrugia & Calvo, 2016). We first tested whether a reduction of the elevated activity of Cdc42 in aged HSCs to the level reported for young HSCs with a specific inhibitor of Cdc42 activity termed CASIN (Florian *et al*, 2012; Leins *et al*, 2018; Grigoryan *et al*, 2018; Liu *et al*, 2019; Amoah *et al*, 2021) might influence distribution of septin 2, 6 or 7 in aged LT-HSCs. Inhibition of Cdc42 activity increased the frequency of aged LT-HSCs polar for septin 7 to the frequency reported for young LT-HSCs (Fig 1C and D) while the distribution of septins 2 and 6 was not affected by CASIN (Fig S1K-M). We also tested whether borgs expressed in LT-HSCs and for which antibodies were available (borg3 (Cdc42ep5) and borg4 (Cdc42ep4) might show co-distribution with septins. Borg4 co-localized with septin 7 in young and aged, CASIN treated LT-HSCs repolarized for septin 7, but not in aged LT-HSCs (Fig 1D), while borg 3 did not co-localize with septin 7 (Fig 1E). In summary, our data support that the activity of Cdc42 regulates the subcellular compartmentalization of HSCs in very specific and targeted manner, due to the fact that the reduction of Cdc42 activity restored the polarity of septin 7 and borg4, but not affecting septin 2, 6 and borg 3 distribution.

We also investigated whether the reduction of Cdc42 activity in aged HSCs by CASIN affected the expression of septins and borgs in LT-HSCs to test for a role of Cdc42 activity in regulating the expression of septins and borgs in addition to controlling their distribution. Interestingly, the level of expression septins 1,2,6,7,8,9 and 11 was increased in CASIN treated aged HSCs, while the expression of septin 8 was decreased and thus also more similar to the level in young LT-HSCs (Fig S1A-G). Similarly, the level of expression of borg 2 and 4 in aged LT-HSCs was more similar to the level in young LT-HSCs upon reduction of Cdc42 activity by CASIN, while the level of borg3 even further increased in aged LT-HSCs upon inhibition of Cdc42 activity (Fig S1H-J). In summary, our data support an influence of Cdc42 activity on the level of expression of distinct borgs and septins, but more importantly on the spatial polar distribution of especially borg4 and septin 7 in LT-HSCs. Interestingly, Borg4 seems to specifically interact with Cdc42 but not to other small RhoGTPases like RhoA or Rac1(Joberty *et al*, 1999, 10; Hirsch *et al*, 2001). On the other hand, septin 7 has a particular role in the septin family due to the fact that it is the only member of its subgroup that cannot be replaced by any other septin in a hexameric or octameric septin assembly (Kremer *et al*, 2005; Tooley *et al*, 2009; Kim *et al*, 2011, 9).

### The Cdc42-borg4-septin 7 interaction in HSCs is regulated by the level of Cdc42 activity

It has been proposed that Cdc42-GTP, but not GDP, binds to borg proteins to ideally position them within the cell while a switch to Cdc42-GDP releases borgs from Cdc42 to allow for interactions with for example septins to stabilize septin networks (Farrugia & Calvo, 2016). We tested such a likely direct physical interaction between Cdc42-borg4 due to their co-distribution in LT-HSCs by a proximity ligation assay (PLA) (Fig 2A). A positive fluorescence signal in a PLA assay indicates a proximity two distinct proteins of at least less than 40 nm between the epitopes which usually equals to physical interaction (Fig S2A). We observed multiple spots of interactions between Cdc42 and borg4 in young HSCs, the level of which was elevated in aged LT-HSCs. Aged LT-HSCs in which Cdc42 activity was adjusted by CASIN showed again a level of reduced interaction similar to young LT-HSCs. This implies that the level of Cdc42 activity regulates the level of the borg4-Cdc42 interaction, with elevated levels of Cdc42-GTP in aged LT-HSCs resulting in enhanced binding of borg4 to Cdc42 (Fig 2A and B). Similarly, we tested for a physical interaction of borg4 with septin 7. There was more interaction in young and aged LT-HSC treated with CASIN, and less in aged LT-HSCs (high Cdc42GTP), implying that borg4 is indeed more likely to be released from Cdc42GDP to then interact with septin 7 (Fig 2B and D). In case of the Cdc42 and borg4 interaction we used Cdc42 knock-out cells as a negative control which indeed delivered only minor background signal (Fig S2B). The attenuation of the elevated activity of Cdc42 in aged HSCs to the level found in young HSCs regulated the extent of the interaction between Cdc42 and borg4 and in turn between borg4 and septin 7. Among the borgs and septins expressed in LT-HSCs, borg4 and septin 7 thus specifically build a critical “effector cascade of Cdc42 activity” to confer outcomes on LT-HSCs (like level of the polar distribution of polarity proteins) in response to changes Cdc42 activity in LT-HSCs.

**Figure 2.**
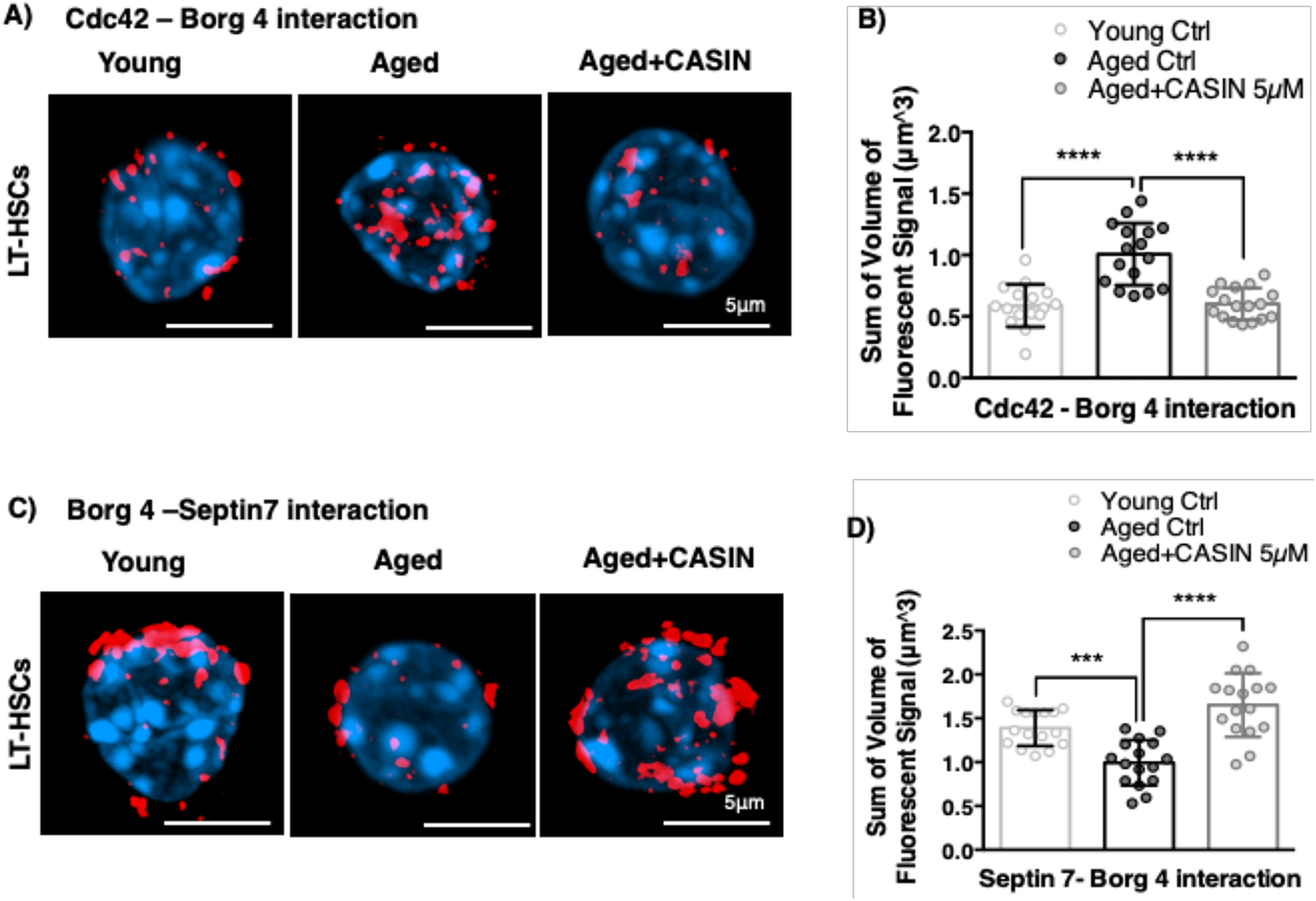
The physical interaction between Cdc42-Borg4-Septin 7 is regulated by the activity of Cdc42. (A&C) Representative immunofluorescence confocal microscopy images of young, aged and aged + CASIN (5µM) treated LT-HSCs for the proximity ligation assay (red signal), testing the extent of close physical interaction of (A) Cdc42 and borg4 and (C) borg4 and septin 7. A red fluorescent signal indicates close proximity of the two proteins tested. Nuclei are stained with DAPI (blue), scale bar= 5µm. (B&D) Quantification of the level of fluorescent signal in young, aged and aged + CASIN (5µM) treated LT-HSCs to quantify the Cdc42-borg4 and borg4-septin 7 proximity the interaction. Data are normalized to the mean fluorescence signal of aged LT-HSCs. 3 biological repeats, n=at least 17 cells per condition, bars = mean±SEM, unpaired t-test, ***P<0.001, ****P<0.0001

### Borg4 or septin 7 regulate the distribution of polarity proteins in HSCs

A general genetic ablation of septin 7 is embryonic lethal, while general genetic ablation of borg4 results in neurological defects (Abbey *et al*, 2016; Menon *et al*, 2014; Ageta-Ishihara *et al*, 2015).To further investigate the role of borg4 and septin 7 for Cdc42 driven phenotypes like polarity in specifically HSCs and hematopoietic cells, we used a vav-1 driven cre-recombinase to delete borg4 or septin 7 in hematopoietic cells of borg4^flox/flox^ or septin 7^flox/flox^ animals (leading to borg4^Δ/Δ^ or septin 7^Δ/Δ^) (Menon *et al*, 2014; Ageta-Ishihara *et al*, 2015). Deletion of borg4 or septin 7 in hematopoietic cells, including LT-HSCs from borg4^Δ/Δ^ or septin 7^Δ/Δ^ animals, was confirmed by PCR, quantitative RT-PCR or immunofluorescence (Fig S3A-D). The absence of borg4 from LT-HSCs resulted in a reduced frequency of LT-HSC polar for the distribution of septin 7 and tubulin but interestingly also for Cdc42 (Fig 3A-C). The absence of septin 7 from LT-HSCs resulted in a reduced frequency of LT-HSC polar for the distribution of borg4, tubulin and, similar to the borg4^Δ/Δ^ LT-HSCs, also for Cdc42 (Fig 3D-F). This strongly implies a strong feedback loop in which changes in the septin 7 localization and thus likely a disturbed septin network will in return influence the localization of Cdc42 as well as borg4. Such a general feedback loop between Cdc42 localization and septins has been previously described in yeast (Okada *et al*, 2013) but not yet for stem cells or even in mammalian cells.

**Figure 3.**
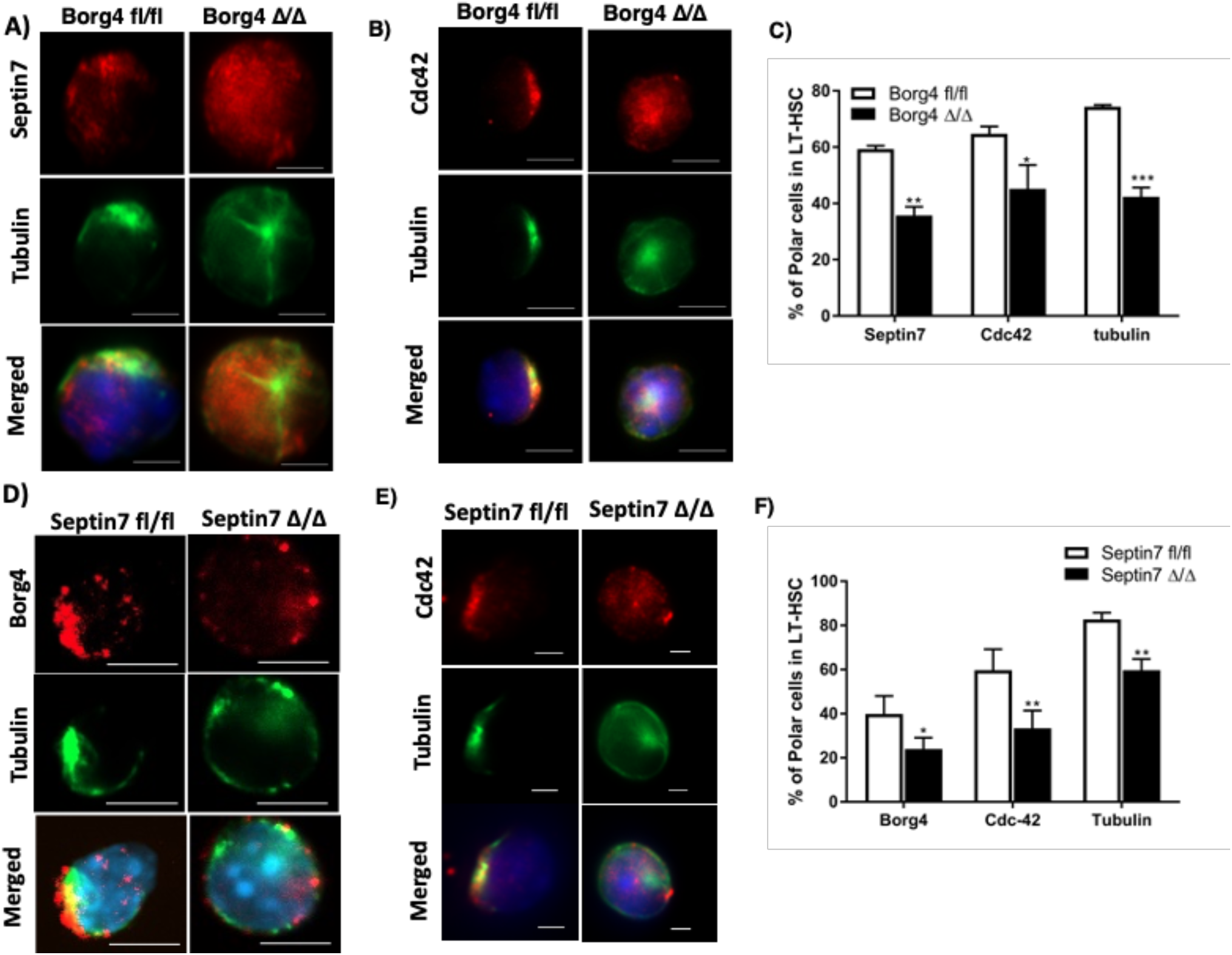
Borg4 or septin 7 regulate the distribution of polarity proteins in HSCs. (A&B) Representative immunofluorescence microscopy images of the distribution of septin 7 (red), Cdc42 (red) and tubulin (green) in LT-HSCs from borg4^fl/fl^ or borg4^Δ/Δ^ mice. Nuclei were stained with DAPI (blue), scale bar= 5 µm. (C) Percentage of cells with a polar distribution of septin 7, Cdc42 or tubulin in borg4^fl/fl^ or borg4^Δ/Δ^ LT-HSCs. At least 3 biological repeats, at least 50 cells were scored per sample. Bars= mean±SEM, one-way ANOVA analysis, **P<0.01, *P<0.05. (D&E) Representative immunofluorescence microscopy images of the distribution of borg4 (red), Cdc42 (red) or tubulin (green) in LT-HSCs from septin 7^fl/fl^ or septin 7^Δ/Δ^ mice. Nuclei were stained with DAPI (blue). scale bar= 5 µm (D) or 2 µm (E). (F) Percentage of cells with a polar distribution of borg4, Cdc42 or tubulin in septin 7^fl/fl^ or septin 7^Δ/Δ^ LT-HSCs. At least 3 biological repeats, at least 30 to 50 cells were scored per sample. Bars= mean±SEM, one-way ANOVA analysis, **P<0.01, *P<0.05.

### HSCs devoid of borg4 or septin 7 show impaired function upon transplantation

The interconnected role of both borg4 and septin 7 for polarity maintenance in LT-HSCs predicts similar changes in the function of HSCs devoid of either borg4 and septin 7 if the level of polarity is linked to HSCs function. We performed analyses of the BM at steady state as well upon competitive transplantation/repopulation. In steady state, the frequency of B- and T- and myeloid cells and the frequency of LT- and ST-HSCs and LMPP was similar in borg4^fl/fl^ and borg4^Δ/Δ^ animals (Fig 4AandB), as well as the frequency of megakaryocyte erythroid progenitors MEPs, common myeloid progenitors (CMPs) and granulocyte macrophage progenitors (GMPs) (Fig S4A). In contrast, the frequency of common lymphoid progenitor cells (CLPs) was significantly increased in borg4^Δ/Δ^ BM (Fig 4C, Fig S4B). Septin 7^Δ/Δ^ animals, similar to borg4^Δ/Δ^ animals, showed frequencies of B-,T- and myeloid cells in BM similar to septin 7^fl/fl^ controls (Fig 4D). Septin 7^Δ/Δ^ animals further presented with a slightly elevated frequency of ST-HSCs but not LT-HSCs, and a reduced frequency of LMPPs (Fig 4E, Fig S4C). Similar to our findings in borg4 ^Δ/Δ^ animals, the frequency of CLPs was also increased in Septin 7^Δ/Δ^ animals (Fig 4F). Next, competitive transplantation/reconstitution experiments with BM cells from fl/fl or Δ/Δ animals were performed and donor chimerism was analyzed up to 21 weeks post-transplantation (Fig 4G). Donor chimerism supported by borg4^Δ/Δ^ cells in both PB and BM was reduced by about 50% in comparison to chimerism supported by borg4^fl/fl^ cells (Fig 4H, Fig S4D), and septin 7^Δ/Δ^ cells almost failed to establish any level of donor chimerism (Fig 4I, Fig S4G). Septin 7^fl/fl^ as well as Septin 7^Δ/Δ^ LSK cells showed a similar efficiency in homing to the BM (Fig S4H), which excludes differential homing to the BM as a cause for the very low level of engraftment of septin 7^Δ/Δ^ cells. Animals reconstituted with borg4^Δ/Δ^ cells presented with a similar frequencies of B-,T- or myeloid cells among donor derived cells in BM (Fig 4J, Fig S4E) or stem and progenitor cells among LSK cells (Fig 4K) or MEPs, CMPs and GMPs in BM (Fig S4F). There was though a significant increase in the frequency of donor-derived CLPs in animals reconstituted with borg4^Δ/Δ^ cells (Fig 4L) which mirrors the findings in steady state hematopoiesis in borg4^Δ/Δ^ animals (Fig 4A-C). Recipients of septin 7 ^Δ/Δ^ cells presented with an increase in the frequency of B220+ B-cells and a decrease in the frequency of myeloid cells in BM (Fig 4M) alongside an increase in the frequency of donor derived LT-HSCs and a reduced LMPP frequency (Fig 4N). The frequency of donor-derived CLPs was significantly increased in recipients of septin 7^Δ/Δ^ cells compared to fl/fl controls (Fig 4O). In summary, these results indicate that in steady state, CLPs frequency is elevated in both borg4^Δ/Δ^ and septin 7^Δ/Δ^ mice, while in general hematopoiesis is, in the case of lack of borg4 or, in the case of lack of septin 7, only slightly affected. Upon competitive transplantation, lack of borg4 and especially septin 7 in HSCs though results in a highly reduced repopulation potential, and in the case of lack of septin 7 also in a skewing of differentiation, with enhanced lymphoid differentiation at the cost of myeloid cells and a higher frequency of LT-HSCs at the cost of the frequency of LMPPs, but similar to steady state, an elevated frequency of CLPs in BM. This data is also consistent with a finding that T-cell specific deletion of septin 7 results in normal T-cell proliferation in vivo, and only impaired T-cell proliferation upon ex vivo expansion.(Mujal *et al*, 2016). Both borg4 and septin 7 are thus necessary for HSC function, especially upon transplantation stress.

**Figure 4.**
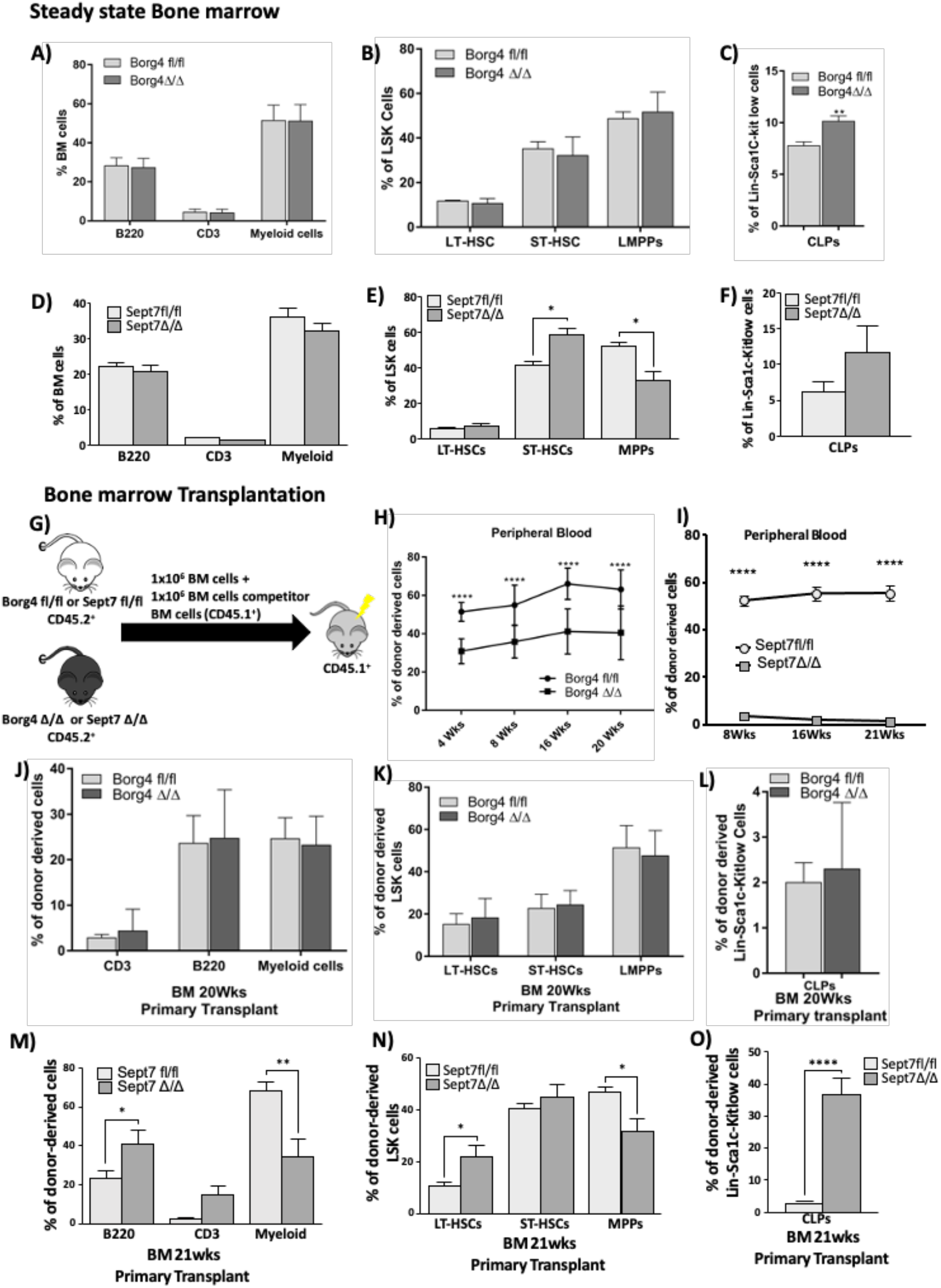
HSCs devoid of borg4 or septin 7 show impaired function upon transplantation. (A-F) Percentage of B-cells (B220 positive cells), T-cells (CD3 positive cells) and myeloid cells (Mac-1^+^ or Gr^+^ positive cells) among BM cells, percentage of LT-HSCs, ST-HSCs and LMPPs in LSK cells and percentage of CLPs in lin^-^Sca1C-kit low cells in borg4^fl/fl^ or borg4^Δ/Δ^ or septin 7^fl/fl^ or septin 7^Δ/Δ^ animals at steady state (n=at least 3). Bars= mean±SD, *P<0.05, Two-way ANOVA analysis. (G) Experimental setup for competitive bone marrow transplantation experiments. (H) Frequency of donor derived cells (borg4^fl/fl^ or borg4^Δ/Δ^) in peripheral blood 4, 8,16 and 20 weeks post transplantation (n=12 per group, Two-way ANOVA analysis). (I) Frequency of donor derived cells (septin 7^fl/fl^ or septin 7^Δ/Δ^) in peripheral blood 4, 8,16 and 20 weeks post transplantation (n=12 per group, Two-way ANOVA analysis). (J-O) Percentage of donor derived (borg4^fl/fl^ or borg4^Δ/Δ^ or septin 7^fl/fl^ or septin 7^Δ/Δ^) B-cells (B220 positive cells), T-cells (CD3 positive cells) and myeloid cells (Mac-1^+^ or Gr^+^ positive cells) among BM cells, percentage of donor derived LT-HSCs, ST-HSCs and LMPPs in LSK cells and percentage of donor derived CLPs in lin^-^Sca1C-kit low cells in animals at steady state (n=12). Bars= mean±SD, ****P<0.0001, ***P<0.001, **P<0.01 and *P<0.05 Two-way ANOVA analysis and 2 tailed unpaired t-test.

### Lack of septin 7 in LT-HSCs impairs HSCs expansion

Lack of septin 7 in HSCs resulted in a severe phenotype upon stress. Septins are known to be essential players in complex processes of compartmentalization, which also include cytokinesis (Kinoshita, 2003). For example, deficiency of septin 7 leads to incomplete cytokinesis in fibroblasts (Menon *et al*, 2014). We therefore tested whether loss of septin 7 might also influence, in addition to polarity, cell proliferation kinetics of LT-HSCs which might contribute then to graft failure. To test proliferation potential, we determined ex vivo single cell division kinetics of LT-HSCs. LT-HSCs from septin 7^Δ/Δ^ mice completed their first and second division an average later compared to HSCs from control septin 7 ^fl/fl^ mice (Fig 5A). Lack of septin 7 in HSCs also resulted in an increase in the frequency of in LT-HSCs dying in the expansion culture (Fig 5B). The number of cells generated from individual septin 7^Δ/Δ^ HSCs 5 days after initiation of the culture was markedly reduced compared to controls (Fig 5C). While these findings do not support a dramatic role of septin 7 for cytokinesis in HSCs, they imply a role for stem cell survival, but more importantly, indicate a rapid loss of HSC and progenitor potential upon cell division as indicated by the small colony size of individual septin 7^Δ/Δ^ HSC.

**Figure 5.**
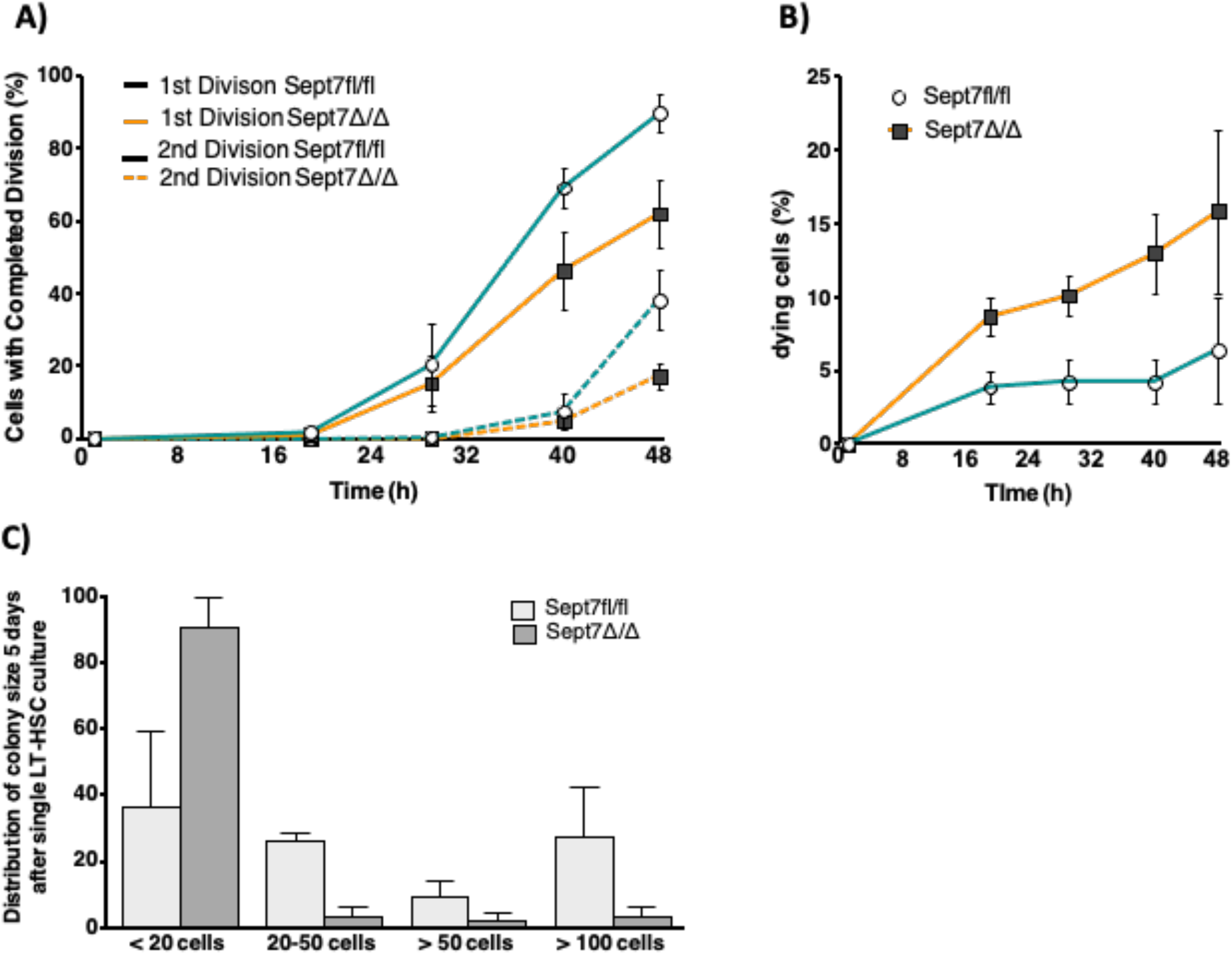
Lack of septin 7 in LT-HSCs impairs HSCs expansion. (A) Percentage of single LT-HSCs from septin 7^fl/fl^ of septin 7^Δ/Δ^ mice that completed first and second division up to 48 hours post initiation of culture. (192 sorted HSCs cells observed). (B) Percentage of single LT-HSCs septin 7^fl/fl^ of septin 7^Δ/Δ^ dying up to 48 hours (192 sorted HSCs observed) (C). Frequency of colonies with distinct cell size 5 days after initiation of culture of individual LT-HSCs from septin 7^fl/fl^ of septin 7^Δ/Δ^ mice (192 sorted HSCs observed).

## Discussion

An increase in the activity of the small RhoGTPase Cdc42 in aged HSCs causes a reduced frequency of polar HSCs and is causative for HSC aging. Polarity within HSCs is tightly linked to the mode of the division. Young (polar) HSCs divide more asymmetrically, while aged (apolar) HSCs divide more symmetrically (Florian *et al*, 2018). Our published data implies that the PAR polarity complex is likely not linked to polarity regulation in HSCs (Florian *et al*, 2012), while we assigned a role of osteopontin or Wnt or Yap-Taz-Scribble signalling in regulating Cdc42 activity and Cdc42 induced HSC polarity (Guidi *et al*, 2017; Florian *et al*, 2013; Althoff *et al*, 2020). We further demonstrated that elevated activity of Cdc42 inhibits the expression of lamin A/C in HSCs, and that lack of this nuclear envelop protein induces loss of epipolarity for H4K16ac (Mejia-Ramirez & Florian, 2020; Grigoryan *et al*, 2018; Mejia-Ramirez *et al*, 2020). However, mechanisms by which Cdc42 activity controls polarity within the cytoplasm of HSCs remained so far not well characterized.

Our data demonstrate that an Cdc42-borg4-septin 7 axis regulates the intracellular distribution of cytoplasmic polarity marker proteins in HSCs in response to changes in Cdc42 activity. Our findings support a model in which Cdc42GTP interacts with borg4, which is then less likely to interact with septin 7 as it is sequestered at Cdc42GTP (Fig 6). On the other hand, if the activity of Cdc42 is reduced and thus the level of Cdc42GTP reduced, borg4 binds to septin 7, which stabilizes septin 7 distribution and by this means likely septin networks to contribute to an orderly compartmentalization within HSCs, as indicated by the elevated frequency of HSCs polar for the distribution of polarity markers.

**Figure 6.**
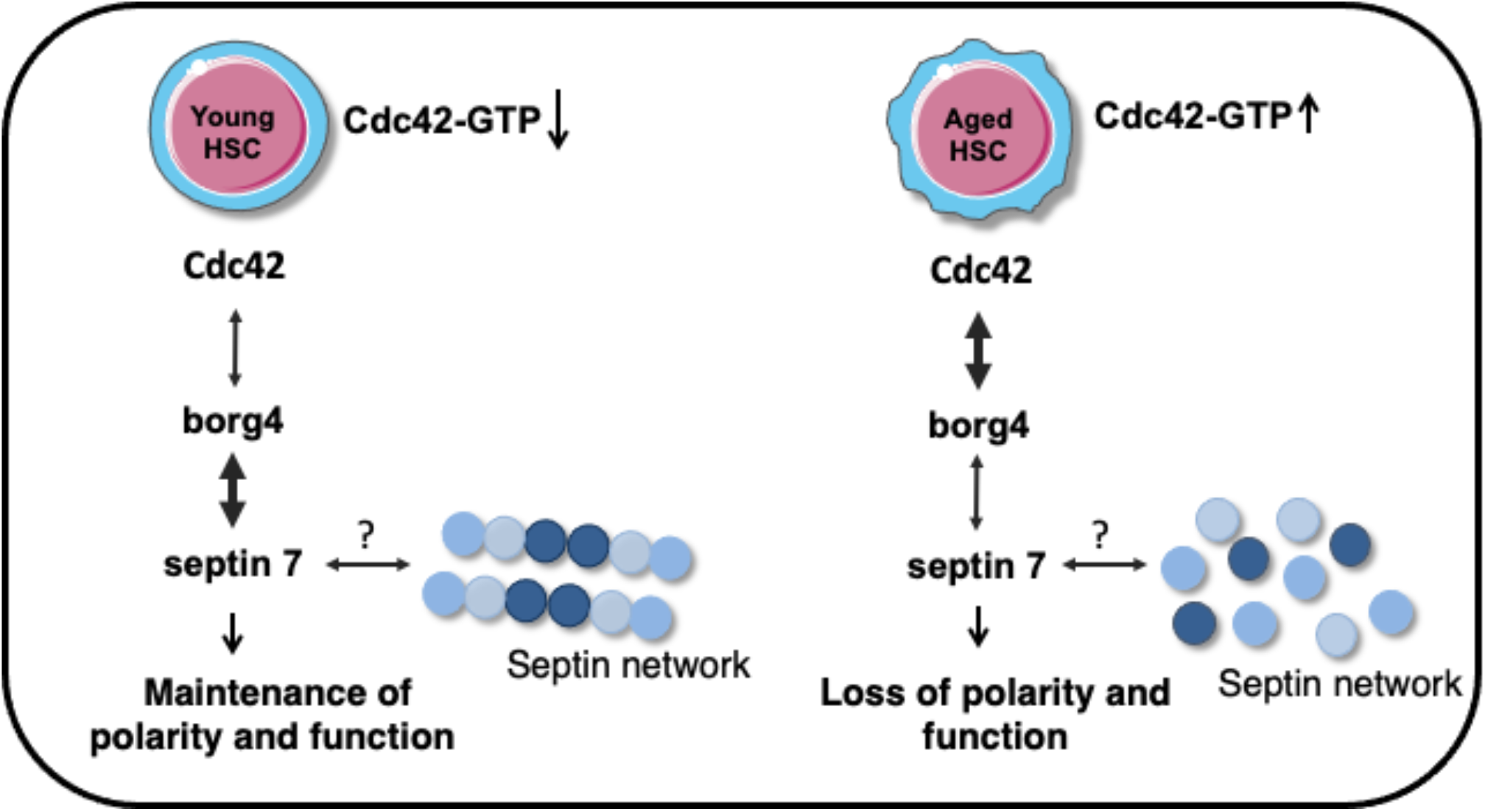
schematic representation of Cdc42-Borg4-Septin 7 axis in the regulation of HSCs polarity and function.

Previous studies reported that ectopic expression of active Cdc42 leads, via borg proteins, to a loss of septin filament assembly in yeast and mammalian cell lines (Joberty *et al*, 2001). This implies that overall similar mechanisms might be at work in yeast and HSCs, with HSCs though specifically relying on borg4 and septin 7 to link Cdc42 activity for the regulation of the position of septins. It is likely that this axis is specific for HSCs, and that thus other cells or cell types might regulate and use borg-septin interactions differentially, with distinct combinations affecting distinct cytoskeletal/nuclear structures. For example borg3 acts via septin 9 to control actomyosin function and cancer cell invasion (Farrugia *et al*, 2020), and a Cdc42-borg2-septin 2 axis activates the DNA damage response in cell lines (Eduardo da Silva *et al*, 2020). It remains to be further investigated whether other types of stem cells will rely on a similar axis, or whether distinct types of stem cells will use type-specific combinations of borgs and septins to link Cdc42 activity to changes in the septin network.

Changes in HSC polarity are linked to changes in HSCs function (Florian *et al*, 2018). Lack of borg4 or septin 7 in HSCs reduced the frequency of polar HSCs and HSCs function was dramatically reduced upon transplantation. It also resulted in an elevated frequency of common lymphoid progenitors (CLPs) in BM while steady-state hematopoiesis was interestingly only marginally affected. Similar changes in the function of HSCs devoid of either borg4 and septin 7 further support an Cdc42-borg4-septin 7 axis in HSCs. The data further imply an important inhibitory role of this borg4-septin 7 axis for the generation of CLPs in vivo, by yet unknown mechanisms. The more severe loss of potential of HSCs that lack septin 7 in comparison to lack of borg4 might be due the structural role of septin 7 in addition to is regulatory role in HSCs, while borg4 having primarily a regulatory role. Another possibility is that other borgs can compensate for the loss of borg4. This might be less likely as we did for example not detect co-localization of borg3 and septin 7 in HSCs (Fig 1E). Although septin 6 is, similar to septin 7, part of the core hexamer complex of septin filaments, lack of septin 6 increased HSCs function as well as the contribution to the B-cell compartment in reconstitution experiments (Senger *et al*, 2017). Together with our finding that core complex septins are not really co-distributed in HSCs (Fig 1D; Fig S1L and M), there is a possibility that only a small fraction of septins within HSCs are, in steady state, actually assembled into filament structures, which though will require further investigations. For example, More recently, it was shown that phosphorylation of septin 1 resulted in impaired cytoskeletal remodeling and thus decreased cell rigidity in HSCs (Ni *et al*, 2019).

While lack of for example Cdc42 in HSCs affects hematopoiesis already at steady state, lack of borg4 or septin 7, effectors of Cdc42 for the regulation of HSCs polarity as shown above, affects HSCs function primarily upon transplantation. Septin 7 is dispensable for the proliferation of B- and T-cells *in vivo* and in the proliferation of splenocytes and myeloid progenitors *in vitro* (Menon *et al*, 2014; Menon & Gaestel, 2015). Interestingly, it was previously shown that successful proliferation of septin 7^Δ/Δ^ T-cells was strictly dependent cell-to-cell contact (Mujal *et al*, 2016). It is thus a possibility that an altered HSC niche interaction that manifests itself only upon transplantation contributes to the lack of proper function of borg4 or sept7^Δ/Δ^ HSCs. This hypothesis is consistent with our finding of reduced colony size of HSCs cultivated ex vivo without proper niche contact (Fig 5C). More recently it was also suggested that HSC contribute only marginally to steady state hematopoiesis, but do so upon transplantation (Busch *et al*, 2015; Dong *et al*, 2020). Another possibility might therefore be that lack of borg4 or septin 7, and in extension, regulation of polarity, might almost exclusively affect HSCs and only to a minor extent more differentiated cells.

Low level of engraftment potential, reduced frequency of HSCs polar for polarity proteins, increased frequency of LT-HSCs upon transplantation and a decreased frequency of LMPPs found in septin 7^Δ/Δ^ animals are also hallmarks of aged HSCs (Florian *et al*, 2012, 42; Adolfsson *et al*, 2005). On the other hand, septin 7^Δ/Δ^ HSCs show elevated contribution to lymphoid differentiation and reduced myeloid differentiation that are not seen in aged HSCs. Lack of borg4 and septin 7, and in extension polarity, thus confer a set of hallmarks of HSC aging, while interestingly lymphoid differentiation is rather enhanced in these animals. Additional studies will be necessary to fully elucidate the role of Cdc42 directed organization of the septin network for the regulation of HSC compartmentalization as well as aging of stem cells.

## Materials and Methods

### Experimental Animals

Young C57BL/6JRj mice (8-12 weeks old) were obtained from Janvier (France), aged C57BL/6J and C57BL/6.SJL *Ptprca Pepcb*/BoyCrl (BoyJ) mice were derived from the internal divisional stock (based in Charles River Laboratories). Septin7 knockout mice (C57BL/6J background) were kindly provided by Matthias Gaestel from Hannover Medical School, Hannover. Borg4 floxed embryos were obtained from RIKEN Bioresource center, Japan. These embryos were utilized to generate initially borg4 floxed mice and further to obtain hematopoietic specific knockout of borg4 gene, borg4 floxed mice crossed with hematopoietic-specific *Vav1-Cre* transgene containing mice. Septin7 and Borg4 mice were confirmed by PCR. The conditional deletion of Septin7 was tested with primer P1: 5′ GGTATAGGGGAC TTTGGGG 3′, primer P2:: 5′ CTTTGCACATATGACTAAGC 3′ and primer P3: 5′ GCTTCTTTTATGTAATCCAGG 3′.(Menon *et al*, 2014). The borg4 conditional deletion was tested with primer a: 5′ TGCTTCAGTACCTTCAGGAC 3′, primer b: 5′ TTCGAGTTCACAGAGCTGGA 3′ and primer c: 5′ TCATAGAGAAGGTGGCAGC 3**′** (Ageta-Ishihara *et al*, 2015, 42). For the Vav-Cre transgene we used the following primers: Cre-FP 5′CTTCTAGGCCTGTACGGAAGTGTT3′ and Cre-RP 5′ CGCGCGCCTGAAGATATAGAAGAT 3′(Croker *et al*, 2004). Cdc42KO mice were housed in the Cincinnati Children’s Hospital Medical Center (CCHMC) animal facility. All mice were housed under pathogen-free environment in the animal barrier facility at the University of Ulm or at CCHMC. Mouse experiments were performed in compliance with the Laws for Welfare of Laboratory Animals and were approved by the Regierungspräsidium Tübingen or by the institutional animal care and use committee at CCHMC.

### Quantitative real time PCR

Gene expression analysis was determined by real-time reverse-transcriptase polymerase chain reaction (RT-PCR) as described previously (Senger *et al*, 2017). Briefly, total RNA was extracted from 20,000 LT-HSCs of young and aged C57BL/6J mice as well as aged LT-HSCs treated with 5µM CASIN using the RNeasy Micro Kit (Qiagen) according to all manufacturer’s instructions. cDNA was prepared and amplified with the Ovation RNA Amplification System V2 (Nu GEN, San Carlos, CA). All real-time polymerase chain reactions were run with Taq MAN Universal PCR Master Mix (Applied Biosystems, Foster City, CA) and with primers from Applied Biosystems on a 7900 HT Fast Real-Time PCR system (Applied Biosystems). Septin1 (Mm00450521_m1), septin2 (Mm00447971_m1), septin3 (Mm00488730_m1), septin4 (Mm00448225_m1), septin5 (Mm01175430_m1), septin6 (Mm00490843_m1), septin7 (Mm00550197_m1), septin8 (Mm01290063_m1), septin9 (Mm01248788_m1), septin10 (Mm00556875_m1), septin11 (Mm01347587_m1),septin12 (Mm01170241_m1), septin14 (Mm01197708_m1), cdc42ep2 (Mm01227921_m1), cdc42ep3 (Mm04208750_m1), cdc42ep5 (Mm00517581_m1), cdc42ep4 (Mm01246312_m1), cdc42ep1 (Mm00840486_m1), Gapdh (Mm99999915_g1).

### Flow cytometry and cell sorting

Peripheral blood and BM cells were analyzed according to stranded procedures and samples were analyzed on LSR Fortessa or LSRII flowcytometry (BD Bioscience). In transplantation to distinguish donors from recipient cells by using Ly45.2 (Clone 104, eBioscience) and Ly45.1(Clone A20, eBioscience) monoclonal antibodies. For PB and BM lineage cell analysis we used the antibodies anti CD3ε (clone 145-2C11), anti B220 (clone RA3-6B2), anti Mac1 (clone M1/70), anti Gr-1(clone RC57BL/6-8C5), all the antibodies from eBiosciences. For early hematopoiesis analysis mononuclear cells were isolated by low density centrifugation (Histopaque 1083, Sigma-Aldrich; Histopaque 1.084, GE Health care) and stained with lineage cocktail antibodies. After that lineage depletion by magnetic separation using Dynabeads (Invitrogen). Lineage depleted cells were stained with anti Sca-1, PE-Cy7 conjugated (clone D7), anti-c-Kit, APC conjugated (clone 2B8), anti-CD34, FITC conjugated (clone RAM34), anti-Flk2, PE conjugated (clone A2F10), anti-IL7R (clone A7R34), anti-CD16/CD32 (clone 93), and streptavidin, eFluor-450 conjugated (all antibodies from eBioscience). FACS analyses data (stem and progenitor cells) were plotted as a percentage of long-term hematopoietic stem cells (LT-HSCs, gated as LSK CD34^-/low^Flk2^−^), short-term hematopoietic stem cells (ST-HSCs, gated as LSK CD34^+^Flk2^−^), and multipotent progenitors (LMPPs, gated as LSK CD34^+^Flk2^+^) distribution among donor-derived LSKs (Lin^neg^c-kit^+^sca-1^+^ cells). Lymphoid and myeloid progenitors are plotted as a percentage of common lymphoid progenitors (CLPs: gated as IL7R^+^ population among Lin^neg^ Sca-1^low^c-Kit ^low^ cells), common myeloid progenitors (CMPs, gated as Lin^neg^c-kit^+^CD34^+^CD16/32^-^), megakaryocyte-erythrocyte progenitors (MEPs gated as Lin^neg^c-kit^+^CD34^-^CD16/32^-^) and granulocyte-macrophage progenitors (GMPs gated as Lin^neg^c-kit^+^CD34^+^CD16/32^+^). FACS analyses data (differentiated cells) are plotted as the percentage of B cells (B220+), T cells (CD3+) and myeloid (Gr-1+, Mac-1+ and Gr-1+Mac-1+) cells among donor-derived Ly5.1+cells. To isolate HSCs lineage depletion was performed to enrich for lineage negative cells, which were then stained with the above-mentioned antibodies and LT-HSCs were sorted using a BD FACS Aria III (BD Biosciences).

### CASIN treatment

Aged LT-HSCs were incubated for 12-16hr with CASIN (5µM) at 37^°^ C (5% CO2,3%O2) in growth factors free medium and subsequently analyzed by immunofluorescence studies or analyzed by real-time RT-PCR.

### Immunofluorescence staining

Freshly sorted LT-HSCs were seeded on fibronectin coated glass coverslips. Cells were fixed with BD Cytofix Fixation Buffer (BD Biosciences). After fixation cells were washed with PBS, permeabilized with 0.2% Triton X-100 (Sigma) in PBS for 20 min, and blocked with 10% Donkey Serum (Sigma) for 20 min. Both primary and secondary antibody incubations were performed for 1hr at room temperature. Coverslips were mounted with ProLong Gold Antifade Reagent with or without DAPI (Invitrogen, Molecular Probes). The cells were coimmunostained with an anti-alpha tubulin antibody (Abcam, rat monoclonal ab 6160), Borg4 (Santa Cruz, goat polyclonal) and Borg3 (Santa Cruz, goat polyclonal) were detected with an anti-rat and goat DyLight 488-conjugated antibody or goat Alexa fluor 647 (Jackson Immuno Research), an anti-Cdc42 antibody (Abcam or Millipore rabbit polyclonal), an anti-AcH4K16 antibody (Millipore, rabbit polyclonal), Septin7 (Proteintech, rabbit polyclonal), Septin6(Santa cruz, rabbit polyclonal), Septin2 (Santa cruz or abcam rabbit polyclonal) were detected with an anti-rabbit DyLight594-conjugated or cy3 conjugated antibody (Jackson Immuno Research). Samples were imaged with an Axio Observer Z1 microscope (Zeiss) equipped with a 63X PH objective. Some of the samples were analyzed with Axio scan microscope with 40X objective. Alternatively, samples were analyzed with an LSM710 confocal microscope (Zeiss) equipped with a 63X objective. Primary raw data were imported into the Volocity Software package (Version 6.0, Perkin Elmer) for further processing.

### Proximity ligation assay

Freshly sorted LT-HSCs were seeded on fibronectin coated glass coverslips. Cells were fixed with BD Cytofix Fixation Buffer (BD Biosciences). After fixation cells were gently washed with PBS, permeabilized with 0.2% Triton X-100 (Sigma) in PBS for 20 min, and blocked with 10% Donkey Serum (Sigma) for 20min. To determine the Cdc42-borg4 and Borg4-septin7 interactions the cells were then incubated with primary antibody mixtures Cdc42 (Millipore rabbit polyclonal) and Borg4(Santa Cruz, goat polyclonal) and Septin7 (Proteintech, rabbit polyclonal) at room temperature for 1hr. Diluted PLA probes were mixed and incubated on the cover glass at 37° C for 1 hr and followed by incubation with ligation mix for 30 min at 37° C. The amplification mix was then applied for 100 min at 37° C. The coverslips were mounted on microscope slides with Doulink Mounting Medium with DAPI, and the cells imaged with a LSM710 confocal microscopy. Further analysed raw data by using Volocity Software package (Version 6.0, Perkin Elmer) to get 3D images and fluorescent intensity calculations.

### Competitive transplantation assays

For competitive bone marrow transplantation assays, young (10–16 weeks old) Sept7^fl/fl^ and Septin7^Δ/Δ^ or Borg4 ^fl/fl^ and Borg4 ^Δ/Δ^ mice were used as donors. A total of 1x 10^6^ donor BM cells were mixed with equal amount of bone marrow cells from Ly5.1^+^ BoyJ competitor mice and transplanted into lethally irradiated Ly5.1^+^ BoyJ recipient mice. Peripheral blood chimerism was determined by FACS analysis up to 21 weeks after primary transplantation. BM cells was analyzed 20 to 21 weeks after primary transplantation. For analysis of the function of septin 7 HSCs, at week 21, 2 × 10^6^ bone marrow cells from individual primary recipient mice were injected into individual lethally irradiated secondary recipient Ly5.1^+^ BoyJ mice.

### Homing assay

LSK cells (CD45.2^+^) were sorted by flow cytometry. LSK cells were cultured with CFSE at 37°C for 8 minutes. The reaction was then terminated with 10% FBS at 4°C for 2 min and washed two times with cold PBS. LKS^+^ cells (1×10^6^) were intravenously injected into lethally irradiated (11 Gy) recipient mice (CD45.1^+^). The recipient mice were sacrificed at 16 hours (h) after transplantation. CFSE^+^ cells in BM of recipient mice were analysed by FACS.

### Single cell proliferation assay

Single cell proliferation assay was performed as described previously (Senger *et al*, 2017). Here, Single LT-HSCs were sorted from the bone marrow of Septin7^fl/fl^ and Septin7 ^Δ/Δ^ mice using BD FACS Aria into the wells of Terasaki plate with culturing medium containing IMDM+ 10% FBS added with 100 ng/mL granulocyte colony-stimulating factor (G-CSF), thrombopoietin (TPO), and stem cell factor (SCF). These cells were incubated for 48 hours at 37°C, 5% CO2, and 3% O2. Within the next 48 hours, the plates were read for completed cell divisions and dead cells every 8 hours. For the first division wells was counted with two cells, second division counted with three to four individual cells.

### Statistical analyses

The statistical analysis were performed within GraphPad prism 7.0. All data were plotted as mean with SD or mean with SEM as minimum and maximum points. Student t-test was used to access the significance difference between the means of two groups. One-way ANOVA and two-way ANOVA analysis were used for more than three groups. Bonferroni post-test to compare all pair of dataset was determined when the overall p value was <0.05.

## Acknowledgement

The authors would like to thank the flow cytometry core facility for cell sorting and for Imaging confocal microscopy at Ulm University and the Tierforschungszentrum at the University of Ulm for their support. We thank the RIKEN Bioresource center, Japan for providing borg4floxed embryos. A.G. was supported by the Marie Curie Ageing Network MARRIAGE funded by the EU. K.S was supported by the RTG 1789 CEMMA funded by the DFG. R.K. was supported by the FOR 2674 funded by the DFG.

## Author contribution

RK, KS and HG were involved in experimental planning and interpretation of results. RK, KS performed experiments. AG performed confocal microscopy analysis. TS performed AxioScan microscopy for imaging. KSo performed cell sorting. VS preformed transplantation experiments. KE performed real time PCR experiments. YZ, MCF and HG supported the overall study design. MM and MG provided Septin 7^fl/fl^ mice and reviewed and edited the manuscript. RK and HG wrote and edited the manuscript.

