## supplementary information for "A Cdc42-Borg4-Septin 7 axis regulates HSCs polarity and function"

**Supplementary figure legends:**

**Figure S1.** (A-G) Level of expression of septin1,2,6,7,8,9 and 11 in young, aged and CASIN (5μM) treated aged LT-HSCs. mRNA levels were normalized to the level of expression of GAPDH and shown relative to the level in young LT-HSCs. N=3, Shown are mean±SEM, unpaired t-test, ****P<0.0001, ***P<0.001, **P<0.01, *P<0.05 (H-J) Level of expression of borg2,3,4 in young, aged and CASIN (5μM) treated aged LT-HSCs. mRNA levels were normalized to the level of expression of GAPDH and shown relative to the level in young LT-HSCs. N=3, shown are mean±SEM, unpaired t-test ***P<0.001, ****P<0.0001 (K) Percentage of young, aged and aged LT-HSCs treated with CASIN (5μM) with a polar distribution of septin2 or septin6. At least 3 biological repeats, at least 50 cells were scored per sample. Bars= mean±SEM, one-way ANOVA analysis. (L&M) Representative immunofluorescence microscopy images of the distribution of septin2 (red) or septin6 (red) or tubulin (green) in young, aged and aged LT-HSCs treated with CASIN (5μM). Nuclei are stained with DAPI (blue), scale bar= 5μm.

**Figure S2.** (A) Schematic representation proximity ligation assay (PLA) reaction for Cdc42-Borg4 and Borg4-Septin7 (B) Representative confocal 3D image of LT-HSC from Cdc42 KO mice tested for the interaction of Cdc42 and Borg4 to determine level of background staining in the PLA assay. Nucleus stained with DAPI.

**Figure S3.** (A) Level of borg4 mRNA in low density bone marrow cells isolated from Borg4$^{fl/fl}$ and Borg4$^{\Delta/\Delta}$ mice (n=3) ****P<0.0001, Bars= mean±SD (B) Detection of floxed and $^{\Delta/\Delta}$ alleles and the Cre transgene of borg4$^{fl/fl}$ or borg4$^{\Delta/\Delta}$ (see also Material and Methods). Two sets of primers were used to distinguish borg4$^{fl/fl}$ (with primer a+b: 580 bp) and borg4$^{\Delta/\Delta}$ (primer a+c: 490bp). The cre allele (375bp) was identified as listed in Material and Methods. (C) Representative Immunofluorescence confocal image of septin7 (red) in LT-HSC septin7$^{fl/fl}$ or septin7$^{\Delta/\Delta}$ mice. Nucleus stained with DAPI. (D) Confirmation of deletion of the septin7 allele in hematopoietic cells: Three primers were to confirm the status of the septin7 alleles in septin7$^{fl/fl}$ and septin7$^{\Delta/\Delta}$ animals. The cre allele (375bp) was identified as listed in Material and Methods.

**Figure S4.** (A) Percentage of MEPs, GMPs and CMPs among $Lin^-IL\text{-}R7^-$Ckit+Sca- cells in BM of borg4$^{fl/fl}$ or borg4$^{\Delta/\Delta}$ animals at steady state. (n=at least 3). Bars= mean±SD, Two-way ANOVA analysis. (B) Gating strategy for the identification of CLPs, GMPs, CMPs and MEPs. (C) Gating strategy for the identification of stem and progenitor cells in BM from septin7$^{fl/fl}$ and septin 7$^{\Delta/\Delta}$ mice at steady state. (D) Frequency of donor (borg4$^{fl/fl}$ or borg4$^{\Delta/\Delta}$) derived cells in BM of primary recipients 20 wks post competitive transplant, bars= mean±SD, **P<0.01, unpaired two-tailed t-test. (E-F) Percentage of donor derived (borg4$^{fl/fl}$ or borg4$^{\Delta/\Delta}$) B-cells (B220 positive cells), T-cells (CD3 positive cells) and myeloid cells (Mac-$1^+$ or $Gr^+$ positive cells) among donor derived PB cells and percentage of donor derived MEPs, CMPs or GMPs among $Lin^-IL\text{-}R7^-$Ckit+Sca- cells in primary recipients 20 wks post competitive transplant (n=12). Bars= mean±SD, unpaired two-tailed t-test. (G) Experimental set up for secondary transplantation and frequency of donor derived cells (septin7$^{fl/fl}$ or septin7$^{\Delta/\Delta}$) in peripheral blood 4, 8,16 and 20 weeks post transplantation (n=12 per group, ****P<0.000, two-way ANOVA analysis). (H) Experimental set up for the determination of the homing ability of sorted LSK cells and relative frequency of CFSE labelled septin7$^{\Delta/\Delta}$ LSK cells homing to bone marrow upon transplantation compared homing of septin7$^{fl/fl}$ cells (set to 1), n=3, shown are mean±SD. unpaired two-tailed t-test.

#### Figure S1

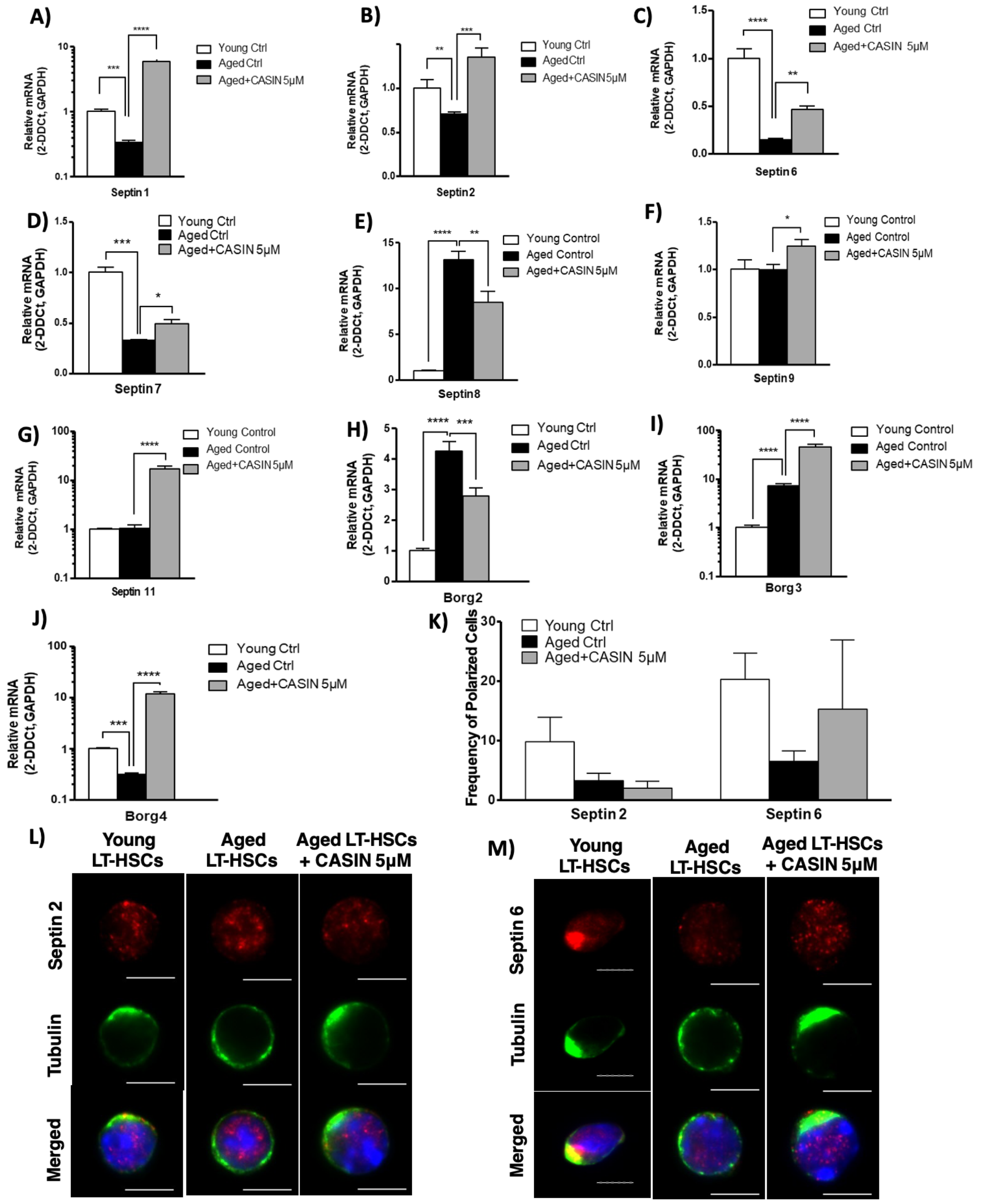

A)
Relative mRNA (2-DDCt, GAPDH)
Septin 1
Young Ctrl
Aged Ctrl
Aged+CASIN 5μM
B)
Septin2
C)
Septin 6
D)
Septin 7
E)
Septin8
Young Control
Aged Control
F)
Septin 9
G)
Septin 11
H)
Borg2
I)
Borg 3
J)
Borg4
K)
Frequency of Polarized Cells
Septin 2
Septin 6
L)
Young LT-HSCs
Aged LT-HSCs
Aged LT-HSCs + CASIN 5μM
Septin 2
Tubulin
Merged
M)
Septin 6

**Figure S2**

A)

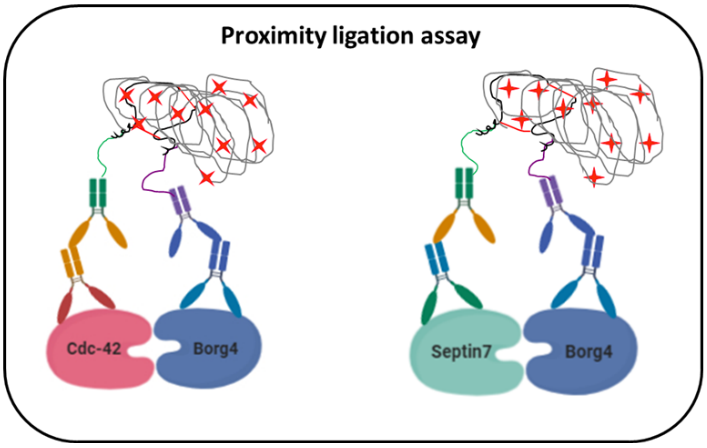
Proximity ligation assay
Cdc-42
Borg4
Septin7
Borg4

B)

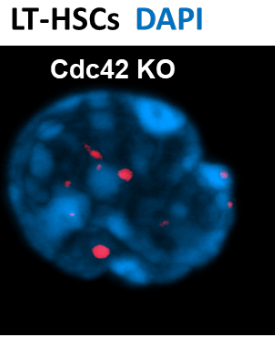
LT-HSCs DAPI
Cdc42 KO

**Figure S3**

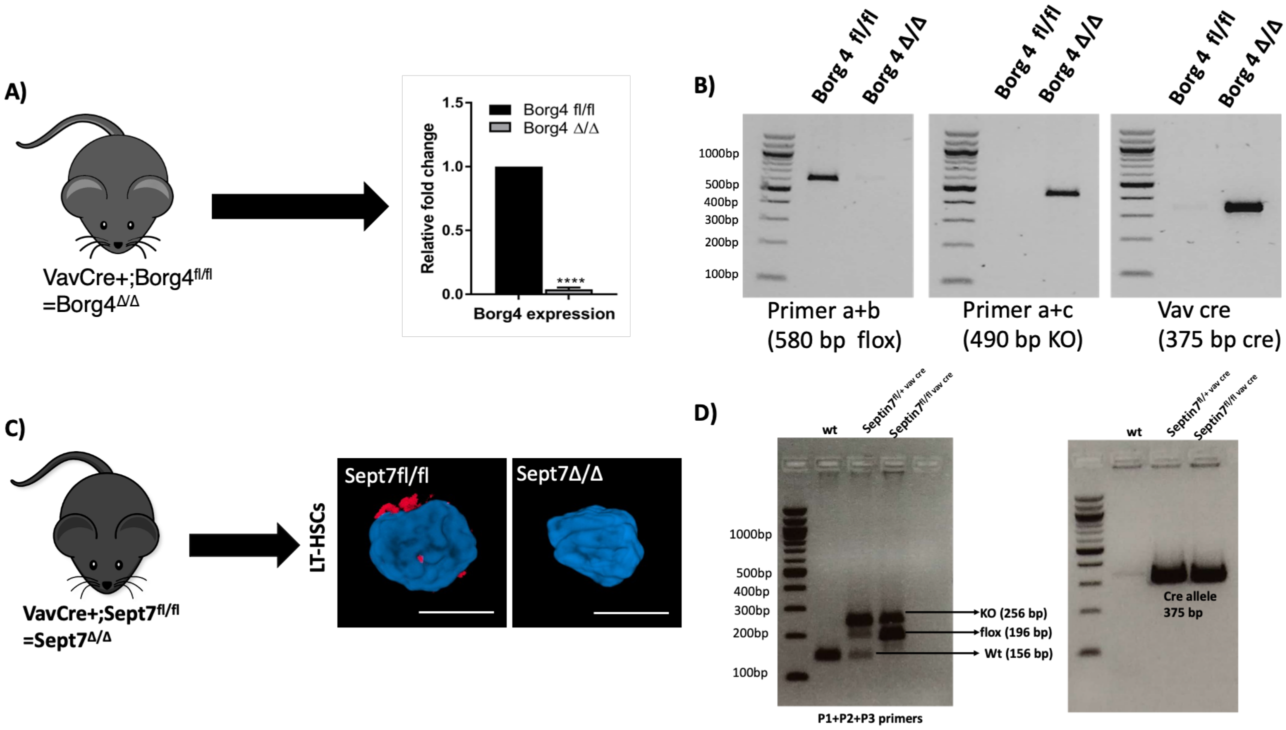

A)
VavCre+;Borg4fl/fl
=Borg4Δ/Δ
1.5
1.0
0.5
0.0
Relative fold change
Borg4 fl/fl
Borg4 Δ/Δ
****
Borg4 expression
B)
Borg 4 fl/fl
Borg 4 Δ/Δ
1000bp
500bp
400bp
300bp
200bp
100bp
Primer a+b
(580 bp flox)
Primer a+c
(490 bp KO)
Vav cre
(375 bp cre)
C)
VavCre+;Sept7fl/fl
=Sept7Δ/Δ
LT-HSCs
Sept7fl/fl
Sept7Δ/Δ
D)
wt
1000bp
500bp
400bp
300bp
200bp
100bp
KO (256 bp)
flox (196 bp)
Wt (156 bp)
P1+P2+P3 primers
Cre allele
375 bp

### Figure S4

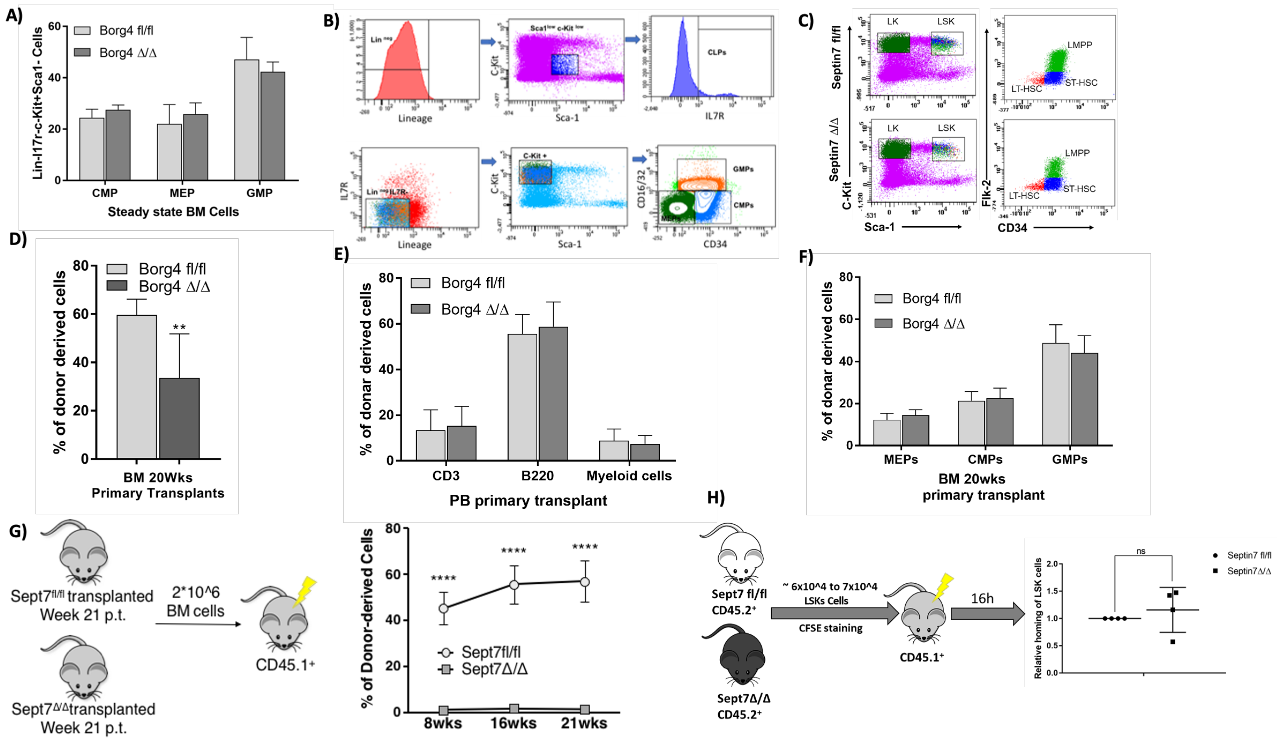

A)
Borg4 fl/fl
Borg4 Δ/Δ
Lin-Il7r-c-Kit+Sca1- Cells
CMP
MEP
GMP
Steady state BM Cells
B)
Lineage
Sca-1
IL7R
CLPs
C-Kit
GMPs
CMPs
CD34
C)
Septin7 fl/fl
Septin7 Δ/Δ
LK
LSK
LMPP
LT-HSC
ST-HSC
C-Kit
Sca-1
Flk-2
CD34
D)
% of donor derived cells
Borg4 fl/fl
Borg4 Δ/Δ
**
BM 20Wks
Primary Transplants
E)
% of donar derived cells
Borg4 fl/fl
Borg4 Δ/Δ
CD3
B220
Myeloid cells
PB primary transplant
F)
% of donar derived cells
Borg4 fl/fl
Borg4 Δ/Δ
MEPs
CMPs
GMPs
BM 20wks
primary transplant
G)
Sept7fl/fl transplanted
Week 21 p.t.
Sept7Δ/Δtransplanted
Week 21 p.t.
2*10^6
BM cells
CD45.1+
% of Donor-derived Cells
****
Sept7fl/fl
Sept7Δ/Δ
8wks
16wks
21wks
H)
Sept7 fl/fl
CD45.2+
Sept7Δ/Δ
CD45.2+
~ 6x10^4 to 7x10^4
LSKs Cells
CFSE staining
CD45.1+
16h
Relative homing of LSK cells
ns
Septin7 fl/fl
Septin7Δ/Δ
